# ChAdOx1 nCoV-19 (AZD1222) vaccine elicits monoclonal antibodies with potent cross-neutralizing activity against SARS-CoV-2 viral variants

**DOI:** 10.1101/2021.09.27.461862

**Authors:** Jeffrey Seow, Carl Graham, Sadie R. Hallett, Thomas Lechmere, Thomas J.A. Maguire, Isabella Huettner, Daniel Cox, Rebekah Roberts, Anele Waters, Christopher C. Ward, Christine Mant, Michael J. Pitcher, Jo Spencer, Julie Fox, Michael H. Malim, Katie J. Doores

## Abstract

Although the antibody response to COVID-19 vaccination has been studied extensively at the polyclonal level using immune sera, little has been reported on the antibody response at the monoclonal level. Here we isolate a panel of 44 anti-SARS-CoV-2 monoclonal antibodies (mAbs) from an individual who received two doses of the ChAdOx1 nCoV-19 (AZD1222) vaccine at a 12-week interval. We show that despite a relatively low serum neutralization titre, mAbs with potent neutralizing activity against the current SARS-CoV-2 variants of concern (B.1.1.7, P.1, B.1.351 and B.1.617.2) were obtained. The vaccine elicited neutralizing mAbs form 8 distinct competition groups and bind epitopes overlapping with neutralizing mAbs elicited following SARS-CoV-2 infection. AZD1222 elicited mAbs are more mutated than mAbs isolated from convalescent donors 1-2 months post infection. Spike reactive IgG+ B cells were still detectable 9-months post boost. These findings give molecular insights into AZD1222 elicited antibody response.

## Introduction

The SARS-CoV-2 encoded Spike glycoprotein is the key target for neutralizing antibodies (nAbs) generated in response to natural infection. The Spike trimer consists of two subunits, S1, that is required for interaction with the ACE-2 receptor on target cells, and S2 that orchestrates membrane fusion. Many monoclonal antibodies (mAbs) have been isolated from SARS-CoV-2 infected individuals allowing identification of key neutralizing epitopes on Spike (Andreano et al., 2021; Barnes et al., 2020; Brouwer et al., 2020; Graham et al., 2021; Piccoli et al., 2020; Robbiani et al., 2020; Rogers et al., 2020; Seydoux et al., 2020; Tortorici et al., 2020). Neutralizing epitopes are present on the receptor binding domain (RBD), the N-terminal domain (NTD) of Spike and S2. RBD-specific nAbs tend to be potently neutralizing and target four epitopes (Barnes et al., 2020; Dejnirattisai et al., 2021; Yuan et al., 2020b), including the receptor binding motif (RBM) which interacts directly with the ACE-2 receptor. Furthermore, several non-overlapping neutralizing epitopes on NTD have been identified which are susceptible to sequence variation in this region (Cerutti et al., 2021; Graham et al., 2021; McCallum et al., 2021; Suryadevara et al., 2021). SARS-CoV-2 infection also generates a large proportion of non-neutralizing antibodies of which the biological function is not fully understood (Anderson et al., 2021; Beaudoin-Bussières, 2021; Li et al., 2021). Combined, studying the antibody response to SARS-CoV-2 infection has generated an antigenic map of the Spike surface (Corti et al., 2021; Dejnirattisai et al., 2021).

Following the emergence of SARS-CoV-2 in the human population, vaccines against COVID-19 have been rapidly developed. Most licenced vaccines use, or encode, a SARS-CoV-2 Spike antigen to elicit both humoral and cellular responses and many have shown remarkable efficacy in Phase III trials (Baden et al., 2021; Polack et al., 2020; Voysey et al., 2021). However, there are concerns that vaccine efficacy could be reduced against newly emerging SARS-CoV-2 variants of concern (VOC), in particular against the alpha (B.1.1.7), beta (B.1.351), gamma (P.1) and delta (B.1.617.2) variants which harbour mutations throughout Spike. Serum neutralizing activity against viral variants has been reported in many double vaccinated individuals, albeit at a reduced potency (Alter et al., 2021; Collier et al., 2021; Edara et al., 2021; Monin et al., 2021; Supasa et al., 2021; Wang et al., 2021d; Zhou and al, 2021). Despite this reduction, real-world data shows current COVID-19 vaccines are still highly effective in preventing severe disease and hospitalizations in locations where SARS-CoV-2 variants of concern are prevalent (Emary et al., 2021; Lopez Bernal et al., 2021; Madhi et al., 2021).

Whilst the antibody response to COVID-19 vaccination has been studied extensively at the polyclonal level using immune sera (Alter et al., 2021; Collier et al., 2021; Dejnirattisai et al., 2021; Edara et al., 2021; Emary et al., 2021; Monin et al., 2021; Supasa et al., 2021; Wall et al., 2021; Wang et al., 2021d; Zhou and al, 2021), little has been reported on the antibody response at the monoclonal level (Amanat et al., 2021; Andreano, 2021; Cho, 2021; Wang et al., 2021d). To address this paucity of information, we isolated a panel of 44 anti-SARS-CoV-2 monoclonal antibodies (mAbs) from an individual (VA14) who had received 2-doses of the AZD1222 (ChAdOx1 nCoV-19) vaccine at a 12-week interval (**Figure 1A**). The AZD1222 vaccine is a replication-defective chimpanzee adenovirus-vectored vaccine expressing the full-length Wuhan SARS-CoV-2 spike glycoprotein gene (Ramasamy et al., 2021; Voysey et al., 2021). Even though low serum neutralization titres (ID_50_ ∼100) were detected in VA14 at 4-months post vaccine booster, nAbs were isolated which displayed potent cross-neutralizing activity against SARS-CoV-2 viral variants of concern (IC_50_ values as low as 0.003 μg/mL). The AZD1222 vaccine elicited NTD- and RBD-specific nAbs that bind epitopes overlapping with nAbs generated following natural infection. Assessment at 9-months post vaccine booster revealed the presence of Spike reactive IgG+ B cells despite undetectable neutralization. These data suggest that although plasma neutralization may be sub-optimal for protection from infection, memory B cells may be sufficient to provide rapid recall responses to protect from serious illness/hospitalizations upon re-infection.

**Figure 1:**
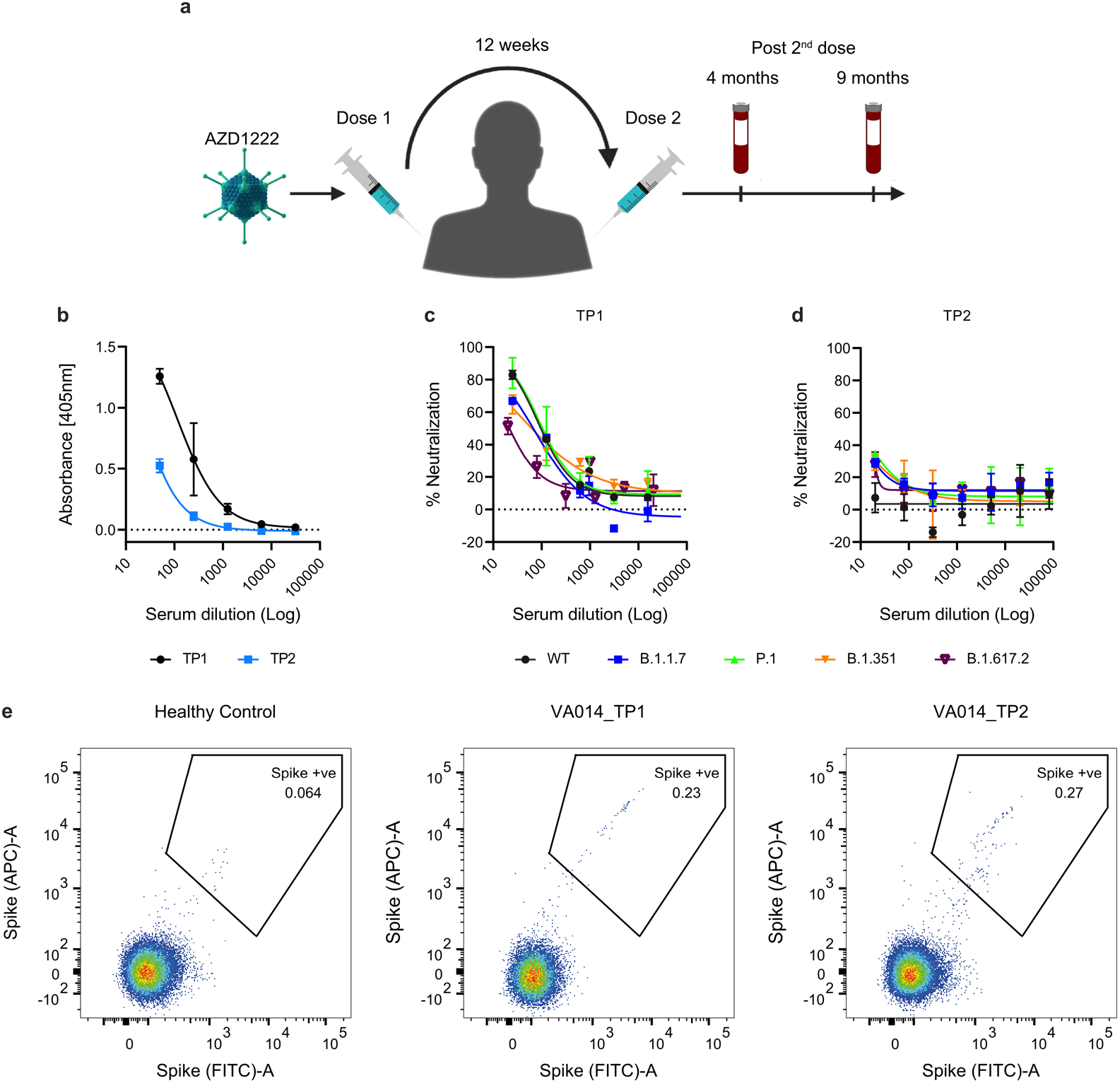
VA14 plasma neutralization and Spike reactive B cells. **A)** Timeline of AZD1222 vaccination and blood sampling for donor VA14. **B)** Plasma IgG binding to Spike at TP1 (4-months post booster) and TP2 (9-months post booster). Plasma neutralizing activity against HIV-1 based virus particles, pseudotyped with the Wuhan, B.1.1.7, P.1, B.1.351 or B.1.617.2 Spike at **C)** TP1 and **D)** TP2. **E)** Fluorescent activated cell sorting (FACS) showing percentage of CD19+IgG+ B Cells binding to SARS-CoV-2 Spike at TP1 and TP2. A healthy control PBMC sample collected prior to the COVID-19 pandemic was used to measure background binding to Spike. The full gating strategy and sorting of RBD specific B cells can be found in **Supplementary Figure 1**.

## Results

### Serum neutralizing activity following AZD1222 vaccination

Plasma and peripheral blood mononuclear cells (PBMC) were isolated from donor VA14 (23 years, white male) at 4-months (timepoint 1, TP1) and 9-months (timepoint 2, TP2) after receiving two doses of the AZD1222 vaccine at a 12-week interval (**Figure 1A**). VA14 reported no previous SARS-CoV-2 infection (based on regular PCR testing), did not have N-specific IgG in their plasma at the time of sampling, and was therefore presumed to be SARS-CoV-2 naïve. Presence of IgG to Spike was determined by ELISA (**Figure 1B**) and a semi-quantitative ELISA measured 0.39 and 0.17 μg/mL of Spike IgG at TP1 and TP2, respectively.

Plasma neutralizing activity was measured using an HIV-1 (human immunodeficiency virus type-1)-based virus particles, pseudotyped with Spikes of SARS-CoV-2 variants of concern, including AZD1222 matched Spike (Wuhan-1, WT), and VOCs B.1.1.7, P.1, B.1.351 and B.1.617.2, and a HeLa cell-line stably expressing the ACE2 receptor (Graham et al., 2021; Seow et al., 2020). Overall, neutralization titres at 4-months post vaccine boost (TP1) were low. ID_50_s of ∼100 were measured against WT and P.1 but were reduced against B.1.1.7, B.1.351 and B.1.351 (**Figure 1C**). Although weak binding to Spike was observed at TP2, neutralization was not detected at a serum dilution of 1:20 (**Figure 1D**).

### Spike reactive B cells detected up to one year following AZD1222 vaccination

Next, we determined the percentage of RBD or Spike reactive IgG expressing B cells at 4- and 9-months post vaccine booster using flow cytometry (**Figure 1E and Figure S1A-D**). 0.25% of IgG+ B cells were Spike reactive and 0.06% were RBD reactive at 4-months post vaccine booster. Despite the undetectable neutralization by sera at 9-months post vaccine booster, 0.27% of IgG+ B cells were Spike reactive.

### AZD1222 vaccination elicits antibodies targeting epitopes on NTD, RBD, S2 and Spike

RBD or Spike reactive B cells at 4-months post vaccine booster were sorted into individual wells and the antibody heavy and light chain genes rescued by reverse transcription followed by nested PCR using gene-specific primers (Graham et al., 2021). Variable regions were ligated into IgG1 heavy and light chain expression vectors using Gibson assembly and directly transfected into HEK 293T/17 cells. Crude supernatants containing IgG were used to confirm specificity to Spike and the variable heavy and light regions of Spike reactive mAbs were sequenced. In total, 44 Spike reactive mAbs were isolated from VA14.

Binding to Spike, S1, RBD, NTD and S2 was determined by ELISA and used to identify the domain-specificity of each mAb (**Figure 2A**). Of the 40 mAbs isolated using the stabilized Spike sorting antigen, 45% (18/40) bound RBD, 35% (14/40) bound NTD, 17.5% (7/40) bound S2 and 2.5% (1/40) bound Spike only (**Figure 2B**). A further four RBD specific mAbs were isolated using the RBD sorting probe. A similar distribution between mAbs targeting RBD, NTD and S2 was seen for mAbs isolated from convalescent donors 6-8 weeks post onset of symptoms (POS) (Graham et al., 2021).

**Figure 2:**
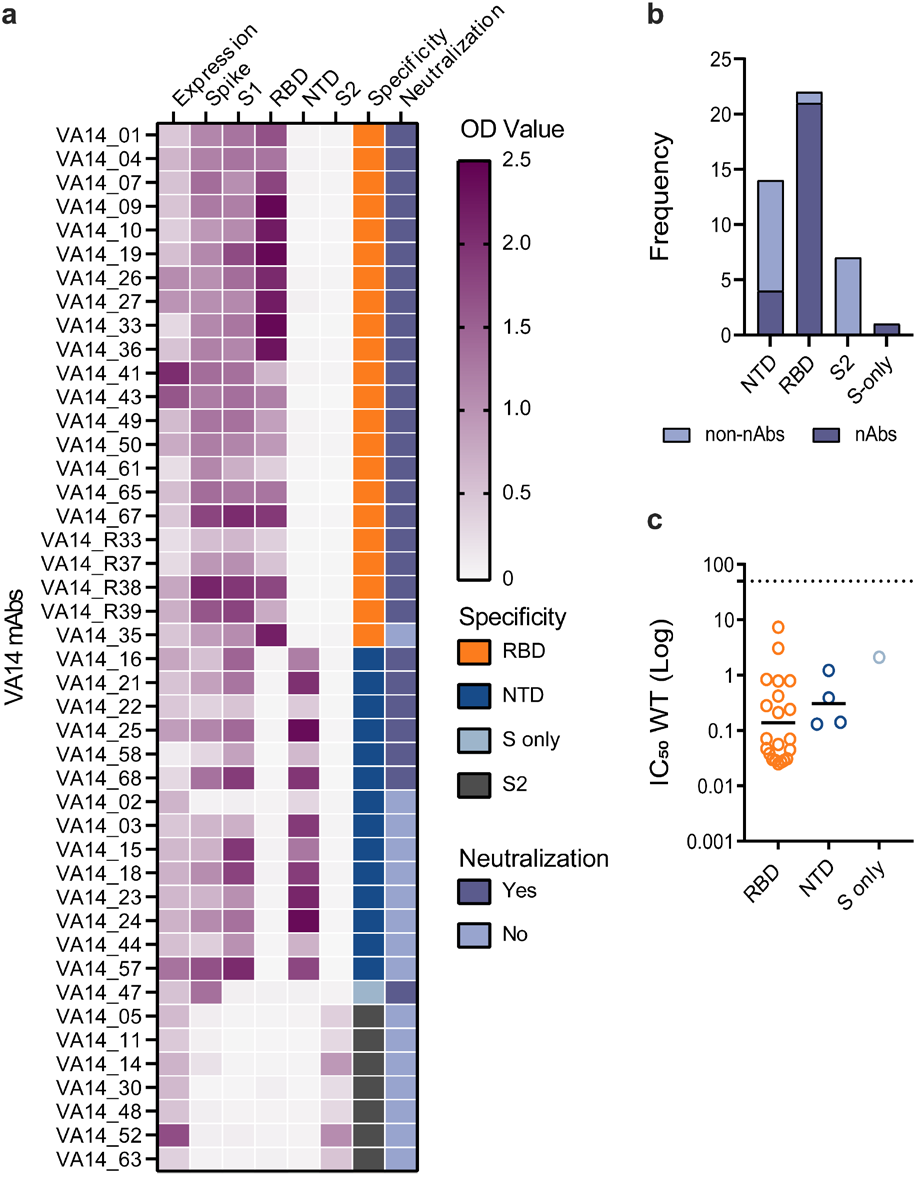
AZD1222 elicits neutralizing and non-neutralizing antibodies targeting RBD, NTD, S1 and S2 domains of Spike. **A)** Heatmap showing IgG expression level and binding to SARS-CoV-2 Spike domains, RBD, NTD, S1 and S2. The figure reports OD values from a single experiment (range 0–2.5) for undiluted supernatant from small scale transfection of 44 cloned mAbs. Antigen binding was considered positive when OD at 405 nm was >0.2 after background was subtracted. SARS-CoV-2 Spike domain specificity for each antibody is indicated. Neutralization activity was measured against wild-type (Wuhan) pseudotyped virus using either small-scale purified IgG or concentrated supernatant. **B)** Frequency of neutralizing and non-neutralizing antibodies targeting either RBD, NTD, S-only or S2. Graph only includes mAbs isolated using Spike as antigen-bait for B cell sorting. **C)** Neutralization potency (IC_50_) against wild-type (Wuhan) pseudotyped virus for mAbs targeting either RBD, NTD or non-S1. The black line represents the geometric mean IC_50_. Related to **Supplementary Figure 2**.

### AZD1222 vaccination elicits neutralizing and non-neutralizing antibodies against epitopes across the full Spike

Neutralizing activity of mAbs was initially measured using HIV-1 virus particles pseudotyped with SARS-CoV-2 Spike encoded by the AZD1222 vaccine. Twenty six of 44 mAbs (59.1%) displayed neutralizing activity of which 21/26 (80.8%) were RBD-specific, 4/26 (15.5%) were NTD-specific and 1/26 (3.8%) only bound Spike (**Figure 2B**). None of the S2-specific nAbs showed neutralizing activity. 95.5% of RBD-specific mAbs and 38.6% of NTD-specific mAbs had neutralizing activity (**Figure 2B**). Neutralization potency against wild-type Spike ranged from 0.025 – 7.3 μg/mL. As previously reported for natural infection, RBD-specific nAbs had a lower geometric mean IC_50_ compared to NTD-specific nAbs (**Figure 2C**) (Graham et al., 2021; Liu et al., 2020).

### AZD1222 elicited mAbs are more highly mutated than mAbs from natural infection

The heavy and light chain variable regions of Spike reactive mAbs were sequenced and the germline usage and level of somatic hypermutation (SHM) determined using IMGT (Brochet et al., 2008). An average 4.9% and 2.8% divergence from V_H_ and V_L_ germlines was observed at the nucleotide level for AZD1222 elicited mAbs (**Figure 3A**), which is higher than mAbs isolated in our previous study from convalescent individuals 3-8 weeks post onset of symptoms (1.9% and 1.4% for V_H_ and V_L_ respectively) (Graham et al., 2021). Three pairs of related clones were identified (**Figure S2A**).

**Figure 3:**
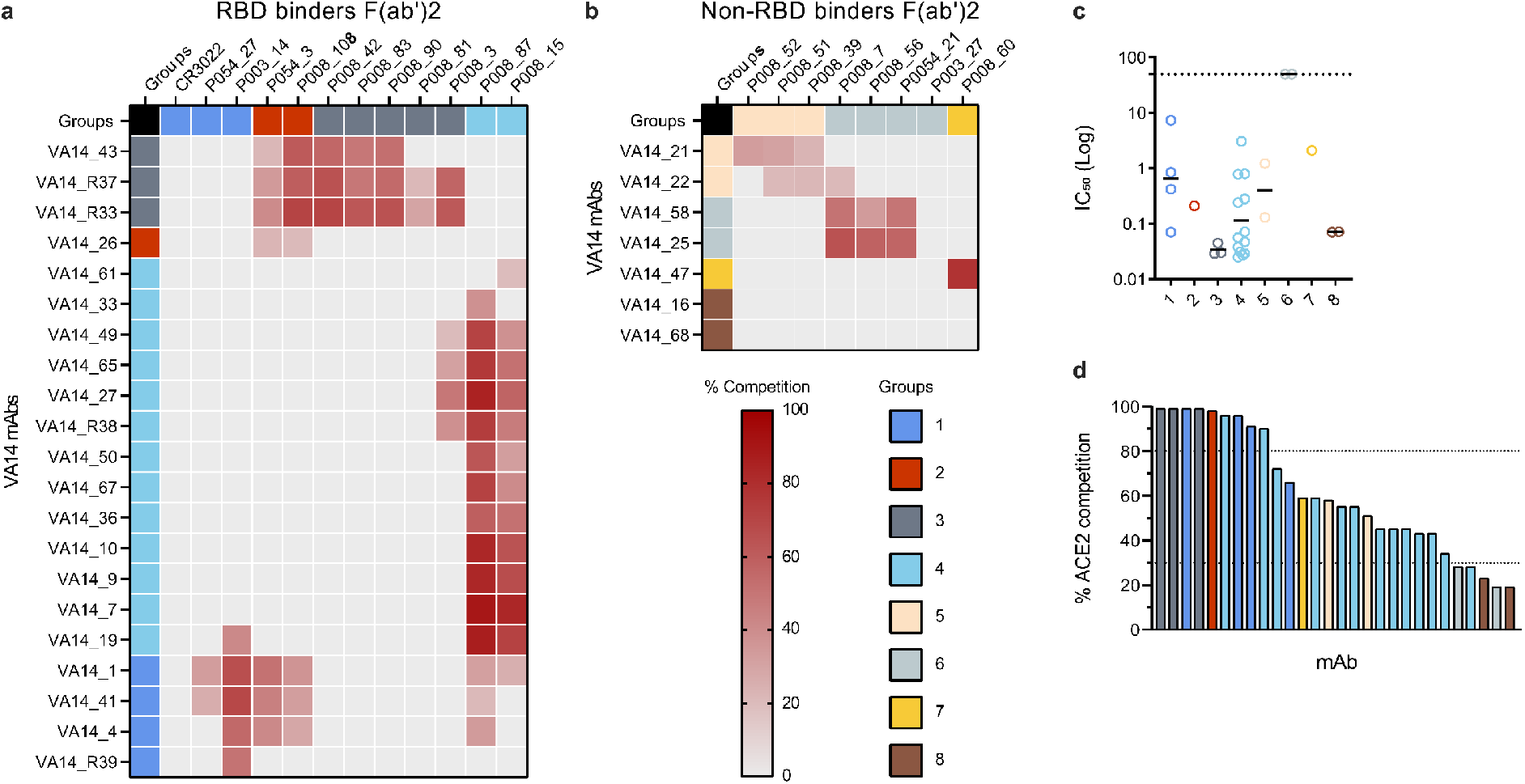
AZD1222 elicited monoclonal antibodies are more mutated than those elicited following SARS-CoV-2 infection. **A)** Truncated violin plot showing the percentage nucleotide mutation compared to germline for the VH and VL genes of Spike-reactive mAbs isolated following AZD1222. Divergence from germline (based on amino acid alignments) for **B)** VH and **C)** VL genes for Spike reactive mAbs from natural infection, AZD1222 vaccination and IgG BCRs from SARS-CoV-2 naïve individuals (Siu, 2021). D’Agostino & Pearson tests was performed to determine normality. Based on the result a Kruskal-Wallis test with Dunn’s multiple comparison post hoc test was performed. *p<0.0332, **p<0.0021, ***p<0.0002 and ****<0.0001. Graph showing the relative abundance of **D)** VH and **E)** VL genes in mAbs elicited from AZD1222 vaccination compared to SARS-CoV-2 infection mAbs (Raybould et al., 2021) and IgG BCRs from SARS-CoV-2 naïve individuals (Siu, 2021). A two-sided binomial test was used to compare the frequency distributions. *p<0.0332, **p<0.0021, ***p<0.0002 and ****<0.0001. Related to **Supplementary Figure 2**.

Germline gene usage and divergence from germline of both neutralizing and non-neutralizing AZD1222 mAbs were compared to a database of SARS-CoV-2 specific mAbs isolated from convalescent individuals (n = 1292) (Raybould et al., 2021) as well as paired heavy and light chains of IgG B cell receptors (BCR) from blood of CD19+ B cells from healthy individuals representative of circulating IgG expressing B cell repertoire (n = 862) (Siu, 2021). As the SARS-CoV-2 mAb database only included amino acid sequences for some mAbs, divergence from germline was determined at the amino acid level (which correlated well with nucleotide divergence (**Figure S2B**)). AZD1222 elicited mAbs from donor VA14 had a statistically higher amino acid mutation (V_H_ 9.2% and V_L_ 6.1%) compared to mAbs isolated from SARS-CoV-2 convalescent donors (V_H_ 4.2% and V_L_ 3.0%) but had a similar level to B cell receptors from healthy subjects (V_H_ 10.9% and V_L_ 8.0%) (**Figure 3B&C**). Similar differences in mutation levels were observed for both neutralizing and non-neutralizing antibodies (**Figure S2C**).

An enrichment in VH3-30 and VH3-53 germline usage was observed for both SARS-CoV-2 infection and AZD1222 elicited mAbs similar to that seen for mRNA elicited mAbs (Wang et al., 2021d) (**Figure 3D**). 3/21 RBD-specific nAbs used the VH3-53/3-66 germlines which are common amongst nAbs that directly bind the ACE2 binding site on Spike (Barnes et al., 2020; Graham et al., 2021; Kim et al., 2021; Robbiani et al., 2020; Yuan et al., 2020c). An enrichment of VH4-34 and VH4-59 germline use was observed for AZD1222 elicited mAbs only. 11/44 (25.0%) and 8/44 (18.2%) mAbs used VK3-20 and VK1-39 light chains, respectively (**Figure 3E**).

### AZD1222 elicited nAbs bind epitopes overlapping with nAbs generated in response to SARS-CoV-2 infection

To gain insight into the epitopes targeted by the AZD1222 elicited nAbs, competition ELISAs with trimeric Spike and previously characterized nAbs isolated from SARS-CoV-2 infected individuals were performed. The panel of competing antibodies encompassed four RBD-, two NTD- and 1 Spike-only competition groups (Graham et al., 2021) (**Figures 4A-B**). Additionally, the ability of nAbs to inhibit the interaction between Spike and the ACE2 receptor was determined by flow cytometry (**Figure 4D**).

**Figure 4:**
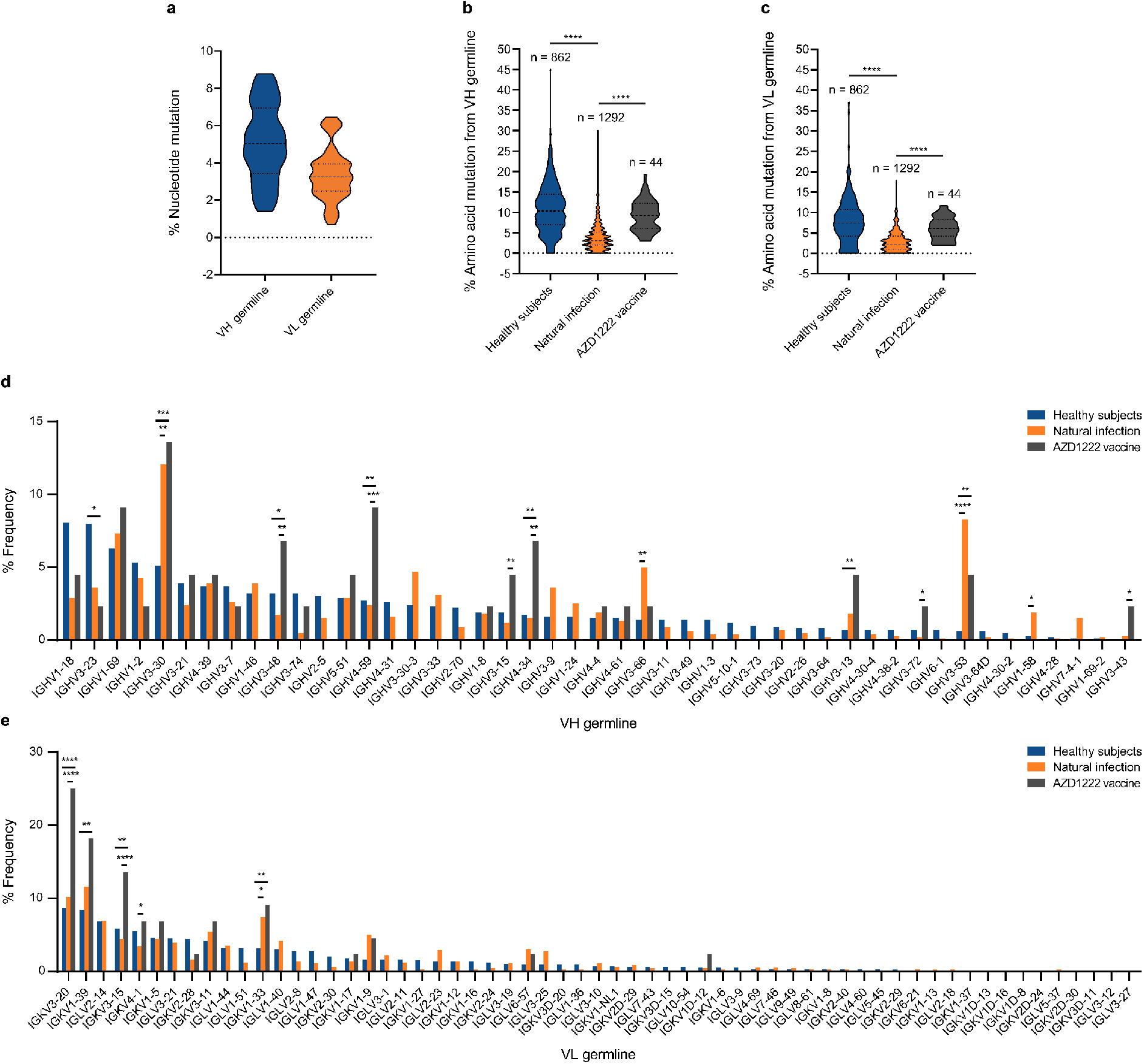
AZD1222 nAbs target epitopes overlapping with nAbs elicited following natural SARS-CoV-2 infection. **A-B)** Competitive binding of AZD1222 and SARS-CoV-2 infection elicited nAbs. Inhibition of IgG binding to SARS-CoV-2 Spike by F(ab)_2_’ fragments was measured. The percentage competition was calculated using the reduction in IgG binding in the presence of F(ab’)_2_ (at 100-molar excess of the IC_80_) as a percentage of the maximum IgG binding in the absence of F(ab’)_2_. Competition was measured between **A)** RBD-specific and **B)** NTD-specific/S-only nAbs. **C)** Neutralization potency (IC_50_) of mAbs targeting either RBD, NTD or non-S1 and/or in competition Groups 1–8 against SARS-CoV-2 WT pseudotyped virus. Competition groups are colour coded according to the key. The black lines represent the geometric mean IC_50_ for each group. IC_50_ values are the average of three independent experiments performed in duplicate. **D)** Ability of nAbs to inhibit the interaction between cell surface ACE2 and soluble SARS-CoV-2 Spike. nAbs (at 600 nM) were pre-incubated with fluorescently labeled Spike before addition to HeLa-ACE2 cells. The percentage reduction in mean fluorescence intensity is reported. Experiments were performed in duplicate. Bars are colour coded based on their competition group.

Four RBD neutralizing antibody classes have been previously identified and characterized (Barnes et al., 2020; Yuan et al., 2020b). nAbs that neutralize by binding to the receptor binding motif (RBM) (equivalent to RBD Class 1) (Barnes et al., 2020; Dejnirattisai et al., 2021; Yuan et al., 2020a) commonly use the VH3-53 or VH3-66 germ lines. As expected, the three VH3-53/VH3-66 VA14 nAbs competed with the Group 3 (RBD Class 1) infection nAbs as well as competing strongly for ACE-2 binding (**Figure 4D**). Group 3 nAbs were most potent at neutralizing the matched vaccine strain (Wuhan-1) (**Figure 4C**).

The majority of RBD-specific nAbs isolated from VA14 (13/20) competed with the Group 4 (RBD Class 3) RBD infection nAbs (**Figure 4A**) and included both potent and modest neutralizing Abs with varying degrees of ACE2 competition (**Figure 4C-D**). Five VA14 nAbs competed with Group 1 (RBD Class 4) RBD infection nAbs and showed a wide range of potencies and levels of ACE2 competition. Only one VA14 nAb (VA14_26) competed with Group 2 (RBD Class 2) RBD infection nAbs which also competed strongly with ACE2.

NTD mAbs formed three competition groups (**Figure 4B**). Non-neutralizing mAbs VA14_25 and VA14_58 competed with NTD Group 6 nAbs including P008_056 which has been shown to bind NTD adjacent to the β-sandwich fold (Rosa et al., 2021b). These two nAbs did not inhibit Spike binding to ACE2 (**Figure 4D**). nAbs VA14_21 and VA14_22 competed with NTD Group 5 nAbs and showed 51-58% inhibition of Spike binding to ACE2. Two NTD nAbs (Group 8) did not compete with any of the infection NTD-specific nAbs or prevent ACE2 binding.

The S-only binding nAb VA14_47 competed with P008_060 (Group 7) (**Figure 4B**), the only other S-only infection nAb, and showed 59% inhibition of Spike binding to ACE2 (**Figure 4D**). P008_060 has been shown to bind a neutralizing epitope on the SD1 domain (*manuscript in preparation*).

### AZD1222 elicited nAbs cross-neutralize SARS-CoV-2 variants of concern

Assessing the cross-neutralizing activity of nAbs isolated from SARS-CoV-2 convalescent donors has revealed that Spike mutations in VOCs selectively hinder neutralizing activity of specific nAb classes (Graham et al., 2021; Wang et al., 2021a; Wang et al., 2021b; Wang et al., 2021c; Wibmer et al., 2021). Therefore, we measured the neutralization potency of AZD1222 elicited nAbs against SARS-CoV-2 variants of concern, including B.1.1.7 (alpha), B.1.351 (beta), B.1.617.2 (delta) and P.1 (gamma) and compared this to nAbs isolated following natural infection (Graham et al., 2021). Spike proteins from these VOCs encode mutations in RBD, NTD and S2 (**Figure 5A**). Some RBD mutations are shared between multiple variants, e.g. B.1.1.7, P.1 and B.1.351 all share an N501Y mutation, and P.1 and B.1.351 share an E484K mutation and a mutation at K417. In contrast, NTD mutations vary considerably between VOCs and include both amino acid mutations and deletions. Although a reduction in neutralization potency was observed for some AZD1222 nAbs, RBD- and NTD-specific nAbs with potent cross-neutralization against all VOCs were identified (**Figure 5B&C**).

**Figure 5:**
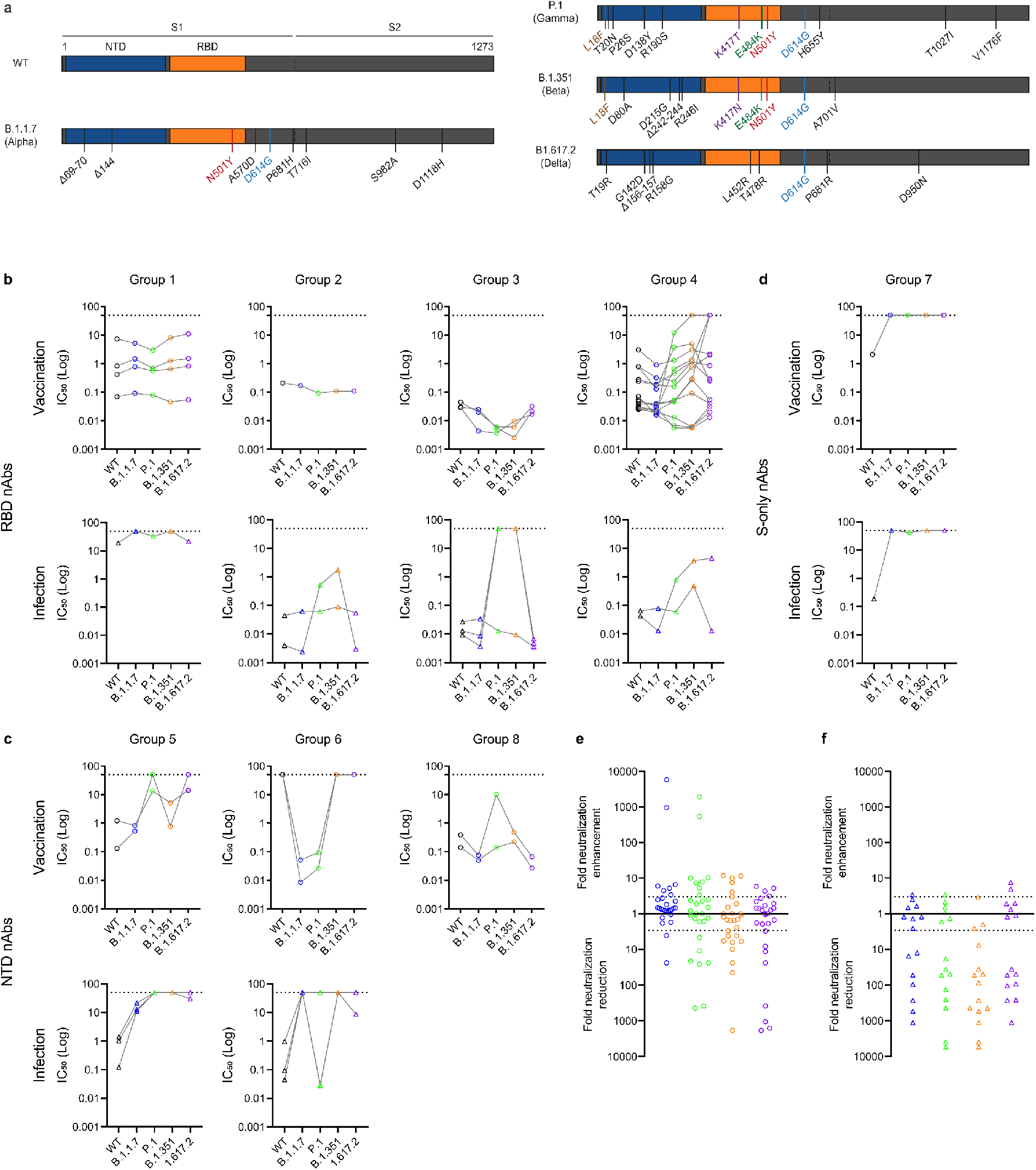
AZD1222 generates nAbs with cross-neutralizing activity against SARS-CoV-2 viral variants. **A)** Schematic showing mutations present in the Spike of SARS-CoV-2 viral variants of concern (B.1.1.7, P.1, B.1.351, B.1.617.2). **B)** Neutralization by RBD-specific nAbs isolated following AZD1222 vaccination or SARS-CoV-2 infection against main variants of concern. nAbs are separated by competition group (Groups 1-4). **C)** Neutralization by NTD-specific nAbs isolated following AZD1222 vaccination or SARS-CoV-2 infection against main variants of concern. nAbs are separated by competition group (Groups 5, 6 and 8). **D)** Neutralization by S-only specific nAbs isolated following AZD1222 vaccination or SARS-CoV-2 infection against main variants of concern. Fold enhancement or reduction in neutralization IC_50_ against VOCs B.1.1.7, P.1, B.1.351, B.1.617.2 compared to the IC_50_ against wild-type for **E)** AZD1222 elicited mAbs and **F)** infection mAbs. The dotted line indicates a 3-fold reduction or enhancement in neutralization. Related to **Supplementary Figure 3 and Supplementary Table 1**.

All Group 3, several Group 4 (VA14_33, VA14_36, VA14R_38) and one Group 1 (VA14R_39) RBD-specific nAbs potently neutralized all five variants at IC_50_s below 0.09 μg/mL (**Figure 5B**). Several nAbs showed enhanced neutralization of VOCs compared to wild-type. Comparing nAbs elicited following infection (Graham et al., 2021) and vaccination, infection nAbs showed a greater sensitivity to Spike mutations in VOCs. Cross-neutralization of nAbs in RBD Groups 1, 2 and 3 was observed for AZD1222 nAbs, whilst some infection nAbs in these competition groups showed greatly reduced neutralization of VOCs P.1 and B.1.351 which both share the E484K mutation. RBD Group 4 mAbs varied in their neutralization of VOCs. 6/13 nAbs showed cross-neutralizing activity. The remaining 7 showed a >3-fold reduction in neutralization against at least one VOC with neutralizing against B.1.351 and B.1.617.2 being most greatly reduced. Despite some RBD nAbs showing a decreased neutralisation against VOCs, binding to variant RBD in ELISA was retained for most nAbs except Group 4 nAbs VA14_19 and VA14_50 (**Figure S3A**) indicating that binding does not always correlate with neutralization.

Considering the geometric mean IC_50_ values, NTD-specific nAbs were most potent at neutralizing the B.1.1.7 VOC. However, the three NTD-competition groups showed differential sensitivities towards the other four SARS-CoV-2 variants (**Figure 5C**). For example, Group 5 NTD nAbs had either reduced or lacked neutralization of P.1 and B.1.617.2, whereas Group 8 NTD nAbs VA14_16 and VA14_68 maintained potent neutralization of B.1.617.2. NTD-specific nAb, VA14_16, had broad reactivity neutralizing all variants with an IC_50_ <0.14 μg/mL and is the only cross-neutralizing NTD-specific nAbs reported thus far (McCallum et al., 2021). Interestingly, two NTD-specific mAbs that had shown no neutralizing activity against WT pseudotyped virus, neutralized both B.1.1.7 and P.1 (**Figure 5C**). The differences in neutralization of VOCs by NTD-specific nAbs were reflected in their binding to S1 of VOCs by ELISA (**Figure S3B**).

The S-only reactive nAbs elicited by vaccination did not neutralize any of the SARS-CoV-2 variants (**Figure 5D**). In contrast, the infection elicited S-only nAb retained modest neutralization against P.1 at concentrations up to 47.7 μg/mL.

Overall, AZD1222 vaccine elicited nAbs showed greater resistance to Spike mutations in variants of concern compared to infection elicited nAbs (**Figure 5E**).

## Discussion

Efficacy of COVID-19 vaccines in the face of SARS-CoV-2 emerging viral variants will be critical for control of the current pandemic. Here we studied the antibody response to the AZD1222 vaccine administered with a 12-week interval at the monoclonal level. The majority of studies examining immune sera from AZD1222 vaccinated individuals have revealed a lower potency against B.1.1.7 (range 2.2 – 9.0-fold) (Dejnirattisai et al., 2021; Emary et al., 2021; Wall et al., 2021), P.1 (2.9-fold) (Dejnirattisai et al., 2021), B.1.351 (range 4.0 – 9.0-fold) (Dejnirattisai et al., 2021; Madhi et al., 2021; Zhou et al., 2021) and B.1.617.2 (range 4.3 – 9.0-fold) (Liu et al., 2021; Wall et al., 2021) compared to neutralization of Wuhan or D614G variants. Although VA14 had a low plasma neutralizing activity (ID_50_ ∼1:100) at 4-months post vaccine booster, 59.1% of Spike reactive mAbs isolated from antigen-reactive B cells had neutralizing activity against the matched vaccine strain, and many of these mAbs displayed potent cross-neutralizing activity against current SARS-CoV-2 VOCs. RBD and NTD were the predominant targets for neutralizing antibodies (80.8% and 15.5% of nAbs, respectively). Importantly, we identified RBD-specific nAbs from each of the four competition groups, and NTD-specific nAbs, that cross-neutralized all VOCs. The polyclonal nature of the nAb response elicited by AZD1222 vaccination will likely help limit full vaccine escape in the face of emerging Spike mutations.

Competition ELISAs revealed that nAbs elicited by AZD1222 target overlapping epitopes of nAbs elicited from natural SARS-CoV-2 infection. However, despite similar antibody footprints, vaccine elicited nAbs from RBD competition Groups 2 and 3 showed greater neutralization breadth than those elicited from natural infection. This was also apparent for some NTD-specific nAbs. This increased neutralization breadth is likely due to the increased divergence from germline in AZD1222 elicited nAbs (isolated 4-months post booster) compared to nAbs isolated following natural infection (isolated 2 – 8 weeks post onset of symptoms) leading to better tolerance of Spike mutations in VOCs. Indeed, several studies have shown that increased somatic hypermutation enhances neutralization breadth against VOCs (de Mattos Barbosa et al., 2021; Gaebler et al., 2021; Goel et al., 2021; Muecksch et al., 2021). Analysis of the antibody-antigen interaction at the molecular level will give further insight into the specific mechanisms of increased neutralization breadth for AZD1222 elicited nAbs.

Although Spike reactive mAbs generated following AZD1222 have not previously been reported, several studies report mAbs isolated following mRNA COVID-19 vaccination (Amanat et al., 2021; Andreano, 2021; Cho, 2021; Wang et al., 2021d). Comparison between epitopes targeted by mRNA and AZD1222 elicited nAbs showed a higher proportion of RBM targeted nAbs following mRNA vaccination (Andreano, 2021; Wang et al., 2021d). A similar enrichment in VH3-53 and VH3-30 germline usage was observed (Andreano, 2021; Wang et al., 2021d). Despite differences in the timing of mAb isolation across reported studies, the AZD1222 mAbs identified had a higher level of SHM compared to mRNA elicited mAbs and showed greater cross-neutralizing activity (Andreano, 2021; Wang et al., 2021d). Possible reasons for these differences include; i) timing of mAb isolation following vaccine booster, ii) timing of vaccine boosters (3-week for mRNA studies vs 12-weeks in this study), iii) a prolonged antigen persistence for ChAdOx vectored Spike, or iv) differences in Spike antigen encoded by each vaccine (in particular, mRNA-1273 (Moderna) and BNT162b2 (Pfizer) vaccines encode Spike with stabilizing mutations and a mutation that prevents S1/S2 cleavage (Jackson et al., 2020; Walsh et al., 2020)). Understanding these factors will be important for optimizing vaccine strategies aimed at eliciting the broadest nAb response.

Plasma was not available to determine the peak neutralizing response in VA14 and therefore the relative decline in neutralization following AZD1222 vaccination. The neutralizing antibody titre was low 4-months post vaccine boost and it is not known if this level would be sufficient to provide sterilizing or near sterilizing immunity. However, the identification of B cells producing antibodies with potent cross-neutralizing activity against non-overlapping epitopes and the presence of Spike+ IgG+ B cells at ∼1 year post vaccine prime suggests that a rapid recall response will likely occur which could be sufficient to protect against severe disease and/or hospitalization in the face of VOCs.

In summary, we show that AZD1222 vaccine administered at a 12-week interval can elicit nAbs with potent cross-neutralizing activity against SARS-CoV-2 VOCs that target non-overlapping epitopes on RBD and NTD. Despite undetectable plasma neutralizing activity, Spike reactive IgG+ B cells are detected up to 1-year following initial vaccine priming. These data provide important insights into long-term immunity and protection to SARS-CoV-2 emerging variants.

### Limitations of study

This study only examines mAbs isolated from one individual and therefore how representative these mAbs are of the humoral immune response against AZD1222 needs to be investigated further.

## EXPERIMENTAL MODEL AND SUBJECT DETAILS

### Ethics

This study used human samples from one donor collected as part of a study entitled “Antibody responses following COVID-19 vaccination”. Ethical approval was obtained from the King’s College London Infectious Diseases Biobank (IBD) (KDJF-110121) under the terms of the IDB’s ethics permission (REC reference: 19/SC/0232) granted by the South Central – Hampshire B Research Ethics Committee in 2019.

### Bacterial strains and cell culture

SARS-CoV-2 pseudotypes were produced by transfection of HEK293T/17 cells and neutralization activity assayed using HeLa cells stably expressing ACE2 (kind gift James E Voss). Small and large scale expression of monoclonal antibodies was performed in HEK293T/17 (ATCC; ATCC^®^ CRL-11268^™^) and 293 Freestyle cells (Thermofisher Scientific), respectively. Bacterial transformations were performed with NEB^®^ Stable Competent *E. coli*.

## METHOD DETAILS

### Protein expression and purification

Recombinant Spike and RBD for ELISA were expressed and purified as previously described (Pickering et al., 2020; Seow et al., 2020). Recombinant S1 (residues 1-530) and NTD (residues 1-310) expression and purification was described in Rosa et al (Rosa et al., 2021a). S2 protein was obtained from SinoBiological (Cat number: 40590-V08B).

For antigen-specific B cell sorting, Spike glycoprotein consisted of the pre-fusion S ectodomain (residues 1–1138) with a GGGG substitution at the furin cleavage site (amino acids 682–685), proline substitutions at amino acid positions 986 and 987, and an N-terminal T4 trimerization domain. RBD consisted of amino acids 331-533. Spike and RBD were cloned into a pHLsec vector containing Avi and 6xHis tags (Aricescu et al., 2006). Biotinylated Spike or RBD were expressed in 1L of HEK293F cells (Invitrogen) at a density of 1.5 × 10^6^ cells/mL. To achieve *in vivo* biotinylation, 480µg of each plasmid was co-transfected with 120µg of BirA (Howarth et al., 2008) and 12mg PEI-Max (1 mg/mL solution, Polysciences) in the presence of 200 µM biotin (final concentration). The supernatant was harvested after 7 days and purified using immobilized metal affinity chromatography and size-exclusion chromatography. Complete biotinylation was confirmed via depletion of protein using avidin beads.

### ELISA (S, RBD, NTD, S2 or S1)

96-well plates (Corning, 3690) were coated with S, S1, NTD, S2 or RBD at 3 μg/mL overnight at 4°C. The plates were washed (5 times with PBS/0.05% Tween-20, PBS-T), blocked with blocking buffer (5% skimmed milk in PBS-T) for 1 h at room temperature. Serial dilutions of plasma, mAb or supernatant in blocking buffer were added and incubated for 2 hr at room temperature. Plates were washed (5 times with PBS-T) and secondary antibody was added and incubated for 1 hr at room temperature. IgM was detected using Goat-anti-human-IgM-HRP (horseradish peroxidase) (1:1,000) (Sigma: A6907) and IgG was detected using Goat-anti-human-Fc-AP (alkaline phosphatase) (1:1,000) (Jackson: 109-055-098). Plates were washed (5 times with PBS-T) and developed with either AP substrate (Sigma) and read at 405 nm (AP) or 1-step TMB (3,3′,5,5′-Tetramethylbenzidine) substrate (Thermo Scientific) and quenched with 0.5 M H_2_S0_4_ before reading at 450 nm (HRP).

### Fab/Fc ELISA

96-well plates (Corning, 3690) were coated with goat anti-human Fc IgG antibody at 3 μg/mL overnight at 4°C. The above protocol was followed. The presence of IgG in supernatants was detected using Goat-anti-human-Fc-AP (alkaline phosphatase) (1:1,000) (Jackson: 109-055-098).

### IgG digestion to generate F(ab’)_2_

IgG were incubated with IdeS (Dixon, 2014) (4 µg of IdeS per 1 mg of IgG) in PBS for 1 hour at 37 °C. The Fc and IdeS A were removed using a mix of Protein A Sepharose® Fast Flow (250 µL per 1 mg digested mAb; GE Healthcare Life Sciences) and Ni Sepharose™ 6 Fast Flow (50 µL per 1 mg digested mAb; GE Healthcare Life Sciences) which were washed twice with PBS before adding to the reaction mixture. After exactly 10 minutes the beads were removed from the F(ab’)_2_-dilution by filtration in Spin-X tube filters (Costar®) and the filtrate was concentrated in Amicon® Ultra Filters (10k, Millipore). Purified F(ab’)_2_ fragments were analysed by SDS-PAGE.

### F(ab’)_2_ and IgG competition ELISA

96-well half area high bind microplates (Corning®) were coated with S-protein at 3μg/mL in PBS overnight at 4 °C. Plates were washed (5 times with PBS/0.05% Tween-20, PBS-T) and blocked with 5% milk in PBS/T for 2 hr at room temperature. Serial dilutions (5-fold) of F(ab’)_2_, starting at 100-molar excess of the IC_80_ of S binding, were added to the plates and incubated for 1 hr at room temperature. Plates were washed (5 times with PBS-T) before competing IgG was added at their IC_80_ of S binding and incubated at room temperature for 1 hr. Plates were washed (5 times with PBS-T) and Goat-anti-human-Fc-AP (alkaline phosphatase) (1:1,000) (Jackson: 109-055-098) was added and incubated for 45 minutes at room temperature. The plates were washed (5 times with PBS-T) and AP substrate (Sigma) was added. Optical density was measured at 405 nm in 5-minute intervals. The percentage competition was calculated as the reduction in IgG binding in the presence of F(ab’)_2_ (at 100-molar excess of the IC_80_) as a percentage of the maximum IgG binding in the absence of F(ab’)_2_. Competition groups were determined using Ward2 clustering (R, Complex Heatmap package (Gu et al., 2016)) for initial analysis and Groups were then arranged by hand according to binding epitopes.

### Semi-quantitative ELISA

In 96-well plates (Corning, 3690), 10 columns were coated with SARS-CoV-2 Spike at 3 µg/mL in PBS, with the remaining 2 columns coated with Goat anti-Human IgG F(ab’)_2_ at 1:1000 dilution, and incubated overnight at 4°C. The plates were washed (5 times with PBS/0.05% Tween-20, PBS-T) and blocked with blocking buffer (5% skimmed milk in PBS-T) for 1 h at room temperature. Serial dilutions of serum and a known concentrations of IgG standard (in blocking buffer) were added to the Spike coated and standard curve columns, respectively. After 2 h incubation at room temperature, plates were washed 5 times with PBS-T. Secondary antibody, goat-anti-human-Fc-AP, was added at 1:1000 dilution in blocking buffer and incubated for 1 h at room temperature. Plates were washed 5 times with PBS-T and developed with AP substrate (Sigma). Absorbance was measured at 405 nm. Antigen-specific serum IgG was quantified by averaging values interpolated from a standard curve of IgG standard using four-parameter logistic regression curve fitting (Rees-Spear et al., 2021).

### SARS-CoV-2 pseudotyped virus preparation

Pseudotyped HIV-1 virus incorporating either the SARS-Cov-2 Wuhan, B.1.1.7, P.1, B.1.351, B.1.617.2 full-length Spike were produced in a 10 cm dish seeded the day prior with 5×10^6^ HEK293T/17 cells in 10 mL of complete Dulbecco’s Modified Eagle’s Medium (DMEM-C, 10% fetal bovine serum (FBS) and 1% Pen/Strep (100 IU/mL penicillin and 100 mg/mL streptomycin)). Cells were transfected using 90 mg of PEI-Max (1 mg/mL, Polysciences) with: 15 µg of HIV-luciferase plasmid, 10 µg of HIV 8.91 gag/pol plasmid (Zufferey et al., 1997) and 5 µg of SARS-CoV-2 spike protein plasmid (Grehan et al., 2015; Thompson et al., 2020). Pseudotyped virus was harvested after 72 hours, filtered through a 0.45mm filter and stored at −80°C until required.

### Neutralization assay with SARS-CoV-2 pseudotyped virus

Neutralization assays were conducted as previously described (Carter et al., 2020; Monin et al., 2021; Seow et al., 2020). Serial dilutions of serum samples (heat inactivated at 56°C for 30mins) or mAbs were prepared with DMEM-C media and incubated with pseudotyped virus for 1-hour at 37°C in 96-well plates. Next, HeLa cells stably expressing the ACE2 receptor (provided by Dr James Voss, Scripps Research, La Jolla, CA) were added (12,500 cells/50µL per well) and the plates were left for 72 hours. The amount of infection was assessed in lysed cells with the Bright-Glo luciferase kit (Promega), using a Victor™ X3 multilabel reader (Perkin Elmer). Measurements were performed in duplicate and duplicates used to calculate the ID_50_.

### Antigen-specific B cell sorting

Fluorescence-activated cell sorting of cryopreserved PBMCs was performed on a BD FACS Melody as previously described (Graham et al., 2021). Sorting baits (SARS-CoV-2 Spike and RBD) was pre-complexed with the streptavidin fluorophore at a 1:4 molar ratio prior to addition to cells. PBMCs were stained with live/dead (fixable Aqua Dead, Thermofisher), anti-CD3-APC/Cy7 (Biolegend), anti-CD8-APC-Cy7 (Biolegend), anti-CD14-BV510 (Biolegend), anti-CD19-PerCP-Cy5.5 (Biolegend), anti-IgM-PE (Biolegend), anti-IgD-Pacific Blue (Biolegend) and anti-IgG-PeCy7 (BD) and Spike-Alexa488 (Thermofisher Scientific, S32354) and Spike-APC (Thermofisher Scientific, S32362) or RBD-Alexa488 and RBD-APC. Live CD3/CD8^-^CD14^-^CD19^+^IgM^-^IgD^-^IgG^+^Spike^+^Spike^+^ or CD3/CD8^-^CD14^-^CD19^+^IgM^-^IgD^-^IgG^+^RBD^+^RBD^+^ cells were sorted using a BD FACS Melody into individual wells containing RNase OUT (Invitrogen), First Strand SuperScript III buffer, DTT and H_2_O (Invitrogen) and RNA was converted into cDNA (SuperScript III Reverse Transcriptase, Invitrogen) using random hexamers (Bioline Reagents Ltd) following the manufacturer’s protocol.

### Full-length antibody cloning and expression

The human Ab variable regions of heavy and kappa/lambda chains were PCR amplified using previously described primers and PCR conditions (Scheid et al., 2009; Tiller et al., 2008; von Boehmer et al., 2016). PCR products were purified and cloned into human-IgG (Heavy, Kappa or Lambda) expression plasmids(von Boehmer et al., 2016) using the Gibson Assembly Master Mix (NEB) following the manufacturer’s protocol. Gibson assembly products were directly transfected into HEK-293T cells and transformed under ampicillin selection. Ab supernatants were harvested 3 days after transfection and IgG expression and Spike-reactivity determined using ELISA. Ab variable regions of heavy-light chain pairs that generated Spike reactive IgG were sequenced by Sanger sequencing.

### IgG expression and purification

Ab heavy and light plasmids were co-transfected at a 1:1 ratio into HEK-293F cells (Thermofisher) using PEI Max (1 mg/mL, Polysciences, Inc.) at a 3:1 ratio (PEI Max:DNA). Ab supernatants were harvested five days following transfection, filtered and purified using protein G affinity chromatography following the manufacturer’s protocol (GE Healthcare).

### ACE2 competition measured by flow cytometry

To prepare the fluorescent probe, Streptavidin-APC (Thermofisher Scientific, S32362) was added to biotinylated SARS-CoV-2 Spike (3.5 times molar excess of Spike) on ice. Additions were staggered over 5 steps with 30 min incubation times between each addition. Purified mAbs were mixed with PE conjugated SARS-CoV-2 S in a molar ratio of 4:1 in FACS buffer (2% FBS in PBS) on ice for 1 h. HeLa-ACE2 and HeLa cells were washed once with PBS and detached using PBS containing 5mM EDTA. Detached cells were washed and resuspended in FACS buffer. 0.5 million HeLa-ACE2 cells were added to each mAb-Spike complex and incubated on ice for 30 m. The cells were washed with PBS and resuspended in 1 mL FACS buffer with 1 μL of LIVE/DEAD Fixable Aqua Dead Cell Stain Kit (Invitrogen). HeLa-ACE2 cells alone and with SARS-CoV-2 Spike only were used as background and positive controls, respectively. The geometric mean fluorescence for PE was measured from the gate of singlet and live cells. The ACE2 binding inhibition percentage was calculated as described previously (Graham et al., 2021; Rogers et al., 2020).

### Monoclonal antibody sequence analysis

The heavy and light chain sequences of SARS-CoV-2 specific mAbs were examined using IMGT/V-QUEST (http://www.imgt.org/IMGT_vquest/vquest) to identify the germline usages, percentage of SHM and CDR region lengths. To remove variation introduced through cloning using mixture of forward primers, 5 amino acids or 15 nucleotides were trimmed from the start and end of the translated variable genes. D’Agostino & Pearson normality test, Kruskal-Wallis test with Dunn’s multiple comparisons post hoc test, Ordinary one-way ANOVA with Tukey’s multiple comparisons post hoc test and two-sided binomial tests) were performed using GraphPad Prism software. Significance defined as p < 0.0332 (*), 0.0021 (**), 0.0002 (***) and >0.0001 (****).

## Acknowledgements

We thank Philip Brouwer, Marit van Gils and Rogier Sanders for the Spike protein construct, Peter Cherepanov for S1 proteins from VOCs, Leo James and Jakub Luptak for the N protein, Wendy Barclay for providing the B.1.617.2 Spike plasmid and James Voss and Deli Huang for providing the Hela-ACE2 cells.

