## Supplementary information for "ChAdOx1 nCoV-19 (AZD1222) vaccine elicits monoclonal antibodies with potent cross-neutralizing activity against SARS-CoV-2 viral variants"

### Supplemental figures:

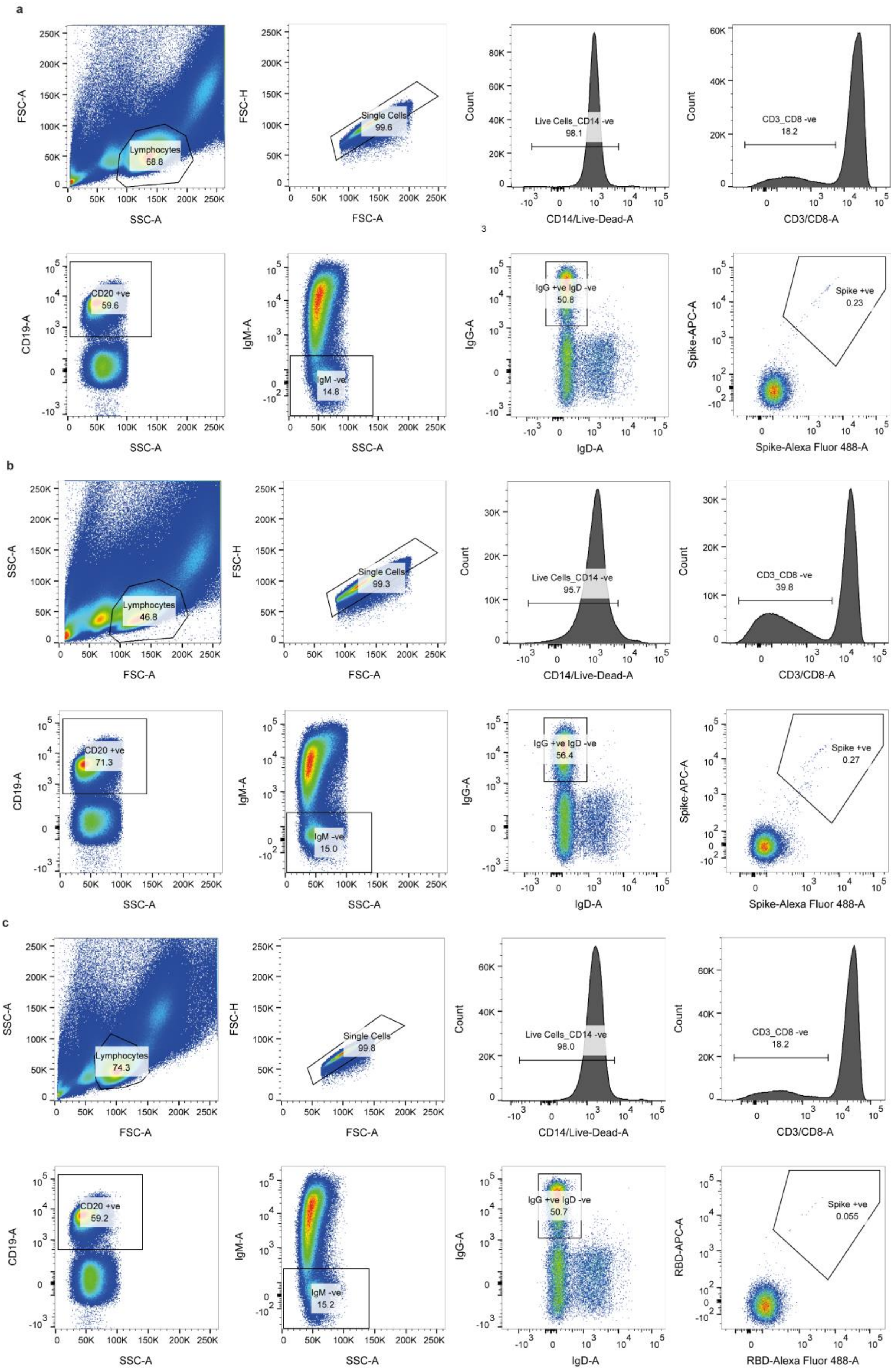

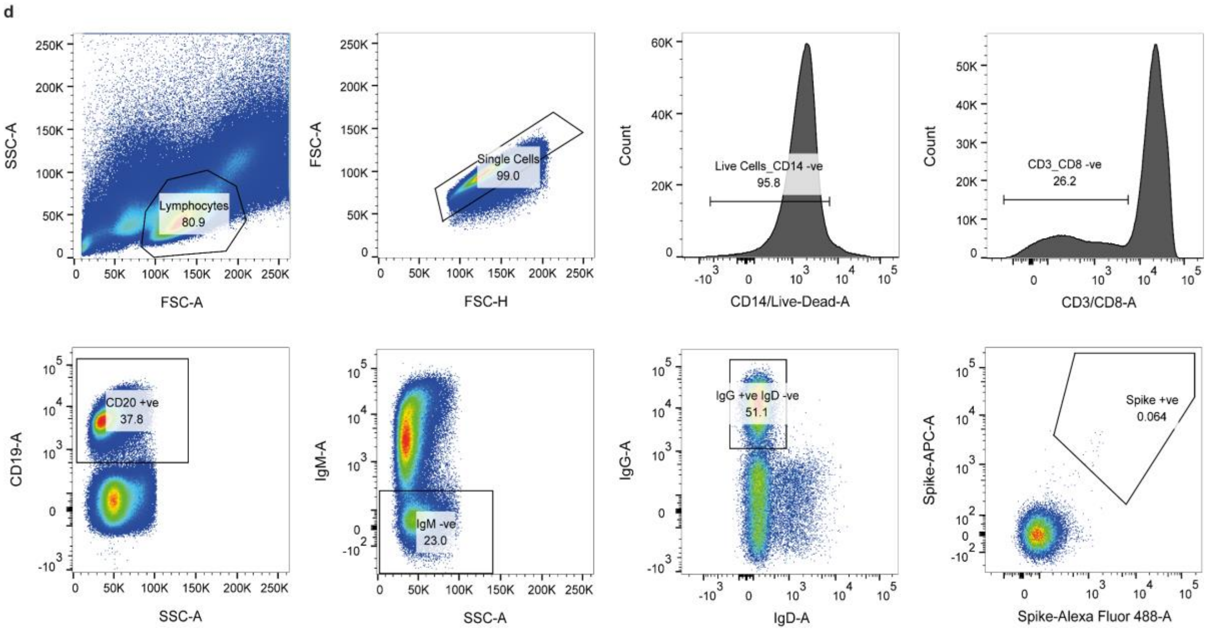


**Supplementary Figure 1: FACS sorting strategy to isolate Spike and RBD reactive mAbs following vaccination with AZD1222.** Sorting of Spike reactive IgG+ B cells from VA14 at **A)** 4-months and **B)** 9-months post vaccine booster. **C)** Sorting of RBD reactive IgG+ B cells from VA14 at 4-months post vaccine booster. **D)** Staining of PBMC collecting pre-COVID-19 from a healthy control.

a

| Name | Protein + Epitope | VH-GENE | JH-GENE | DH-GENE | VL-GENE | JL-GENE | CDRH3 Length | CDRL3 Length | Neutralzation | % Identity Heavy | % Identity Light |
|---|---|---|---|---|---|---|---|---|---|---|---|
| VA014_09 | RBD | IGHV1-18 | IGHJ3 | IGHD3-10 | IGKV3-11 | IGKJ4 | 16 | 8 | Yes | 95.41 | 96.55 |
| VA014_50 | RBD | IGHV1-18 | IGHJ3 | IGHD3-10 | IGKV3-11 | IGKJ4 | 16 | 8 | Yes | | |
| VA014_R39 | RBD | IGHV4-34 | IGHJ6 | IGHD4-17 | IGKV1-5 | IGKJ2 | 18 | 8 | Yes | 87.1 | 94.36 |
| VA014_26 | RBD | IGHV4-34 | IGHJ6 | IGHD4-17 | IGKV1-5 | IGKJ2 | 18 | 8 | Yes | | |
| VA014_27 | RBD | IGHV5-51 | IGHJ3 | IGHD2-2 | IGKV1-39 | IGKJ2 | 23 | 9 | Yes | 95.38 | 95.03 |
| VA014_67 | RBD | IGHV5-51 | IGHJ6* | IGHD2-2 | IGKV1-39 | IGKJ2 | 23 | 9 | Yes | | |

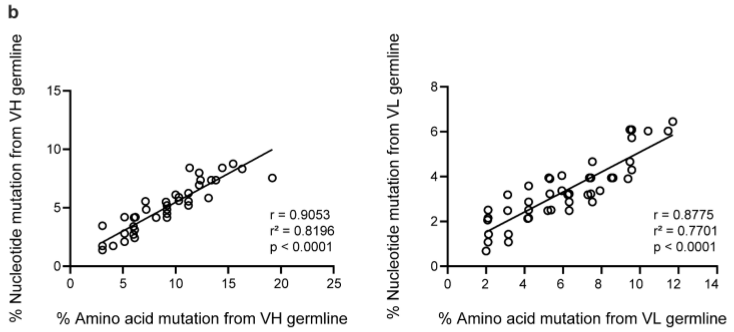


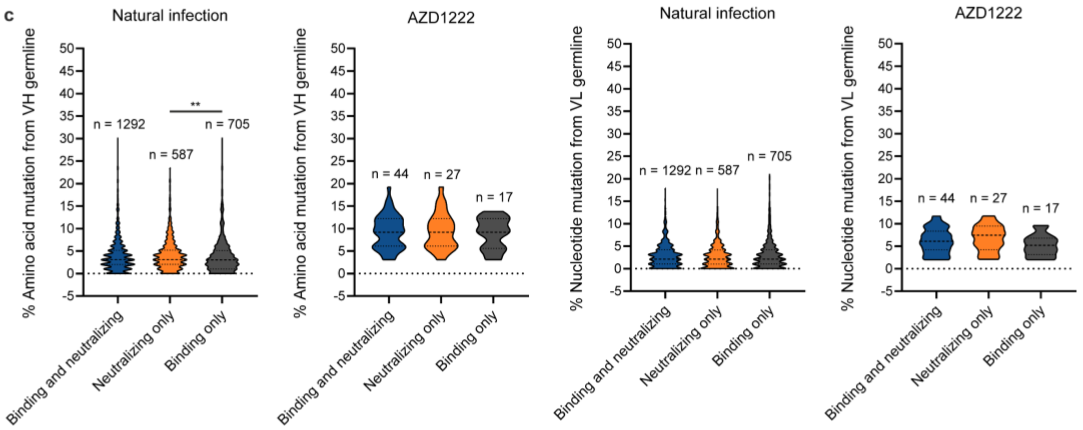


**Supplementary Figure 2: Sequence analysis of AZD1222 elicited mAbs. A)** Clonally related mAbs isolated from VA14. **B)** Correlation between the % nucleotide mutation and % amino acid mutation for VH and VL germline for AZD1222 elicited mAbs (Spearman correlation, two-tailed, r). A linear regression was used to calculate the goodness of fit ($r^2$). **C)** Divergence from germline (based on amino acid alignments) for VH and VL genes for Spike reactive mAbs arising from natural infection and AZD1222 vaccination. Spike reactive mAbs have been separated based on their binding and/or neutralizing properties. D'Agostino & Pearson tests were performed on each dataset to determine normality. Based on the result, either a Kruskal-Wallis test with Dunn's multiple comparison post hoc test or an ordinary one-way ANOVA with Turkey's multiple comparison post hoc test was performed. *p<0.0332, **p<0.0021, ***p<0.0002 and ****<0.0001.

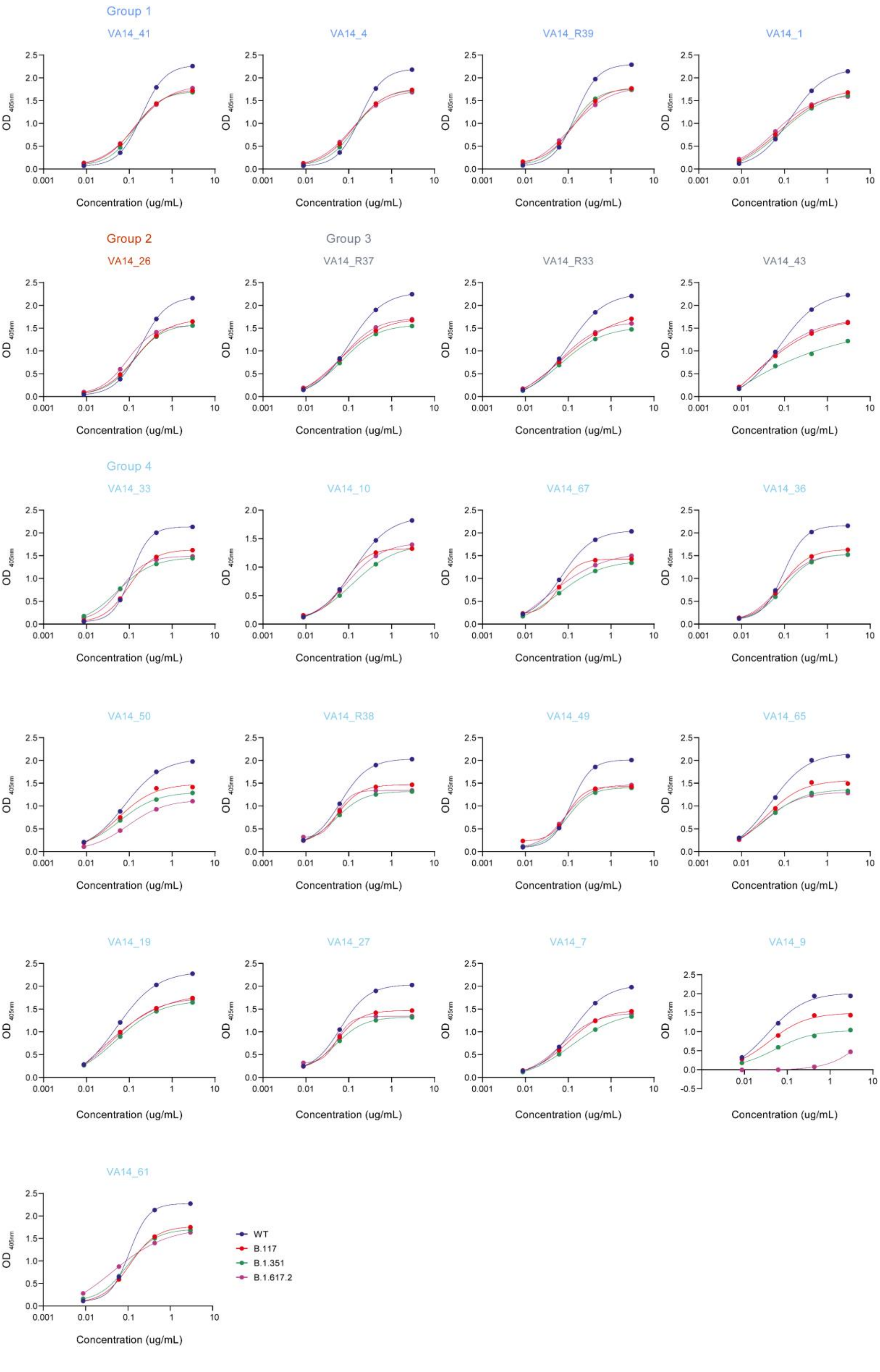
Group 1
VA14_41
VA14_4
VA14_R39
VA14_1
Group 2
VA14_26
Group 3
VA14_R37
VA14_R33
VA14_43
Group 4
VA14_33
VA14_10
VA14_67
VA14_36
VA14_50
VA14_R38
VA14_49
VA14_65
VA14_19
VA14_27
VA14_7
VA14_9
VA14_61
OD 405nm
Concentration (ug/mL)
WT
B.117
B.1.351
B.1.617.2

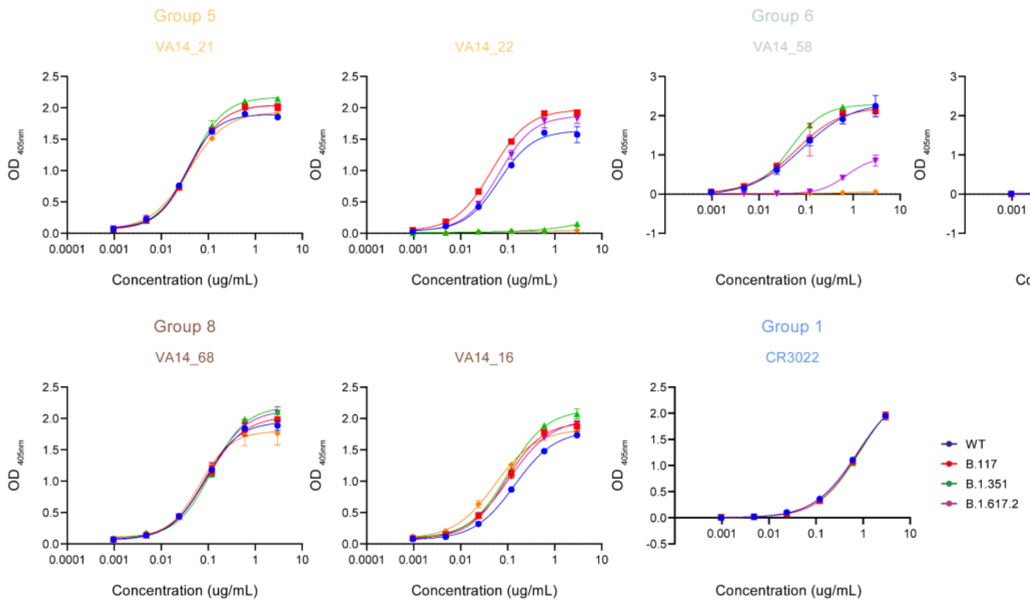


**Supplementary Figure 3: Binding of AZD1222 neutralizing antibodies to RBD or S1 from SARS-CoV-2 variants of concern. A)** Binding of RBD-specific nAbs to recombinant RBD from WT, B.1.1.7, B.1.351 and B.1.617.2 by ELISA. **B)** Binding of NTD-specific nAbs to recombinant S1 from WT, B.1.1.7, B.1.351 and B.1.617.2 by ELISA.

| | Specificity | Competition Group | ACE2 % Competition | WT IC50 | B.1.1.7 IC50 | P.1 IC50 | B.1.351 IC50 | B.1.617.2 IC50 | VH | VL |
|---|---|---|---|---|---|---|---|---|---|---|
| **VA14_R39** | RBD | 1 | 99 | 0.07 | 0.091 | 0.079 | 0.046 | 0.055 | IGHV4-34 | IGKV1-5 |
| **VA14_04** | RBD | 1 | 96 | 0.42 | 0.78 | 0.56 | 0.65 | 0.82 | IGHV3-13 | IGKV1-39 |
| **VA14_41** | RBD | 1 | 91 | 0.84 | 1.46 | 0.69 | 1.26 | 1.52 | IGHV3-30 | IGKV1-39 |
| **VA14_01** | RBD | 1 | 66 | 7.34 | 5.18 | 2.94 | 8.17 | 11.08 | IGHV3-13 | IGKV1-39 |
| **VA14_R37** | RBD | 3 | 99 | 0.03 | 0.025 | 0.0057 | 0.0026 | 0.021 | IGHV3-53 | IGKV3-20 |
| **VA14_R33** | RBD | 3 | 99 | 0.045 | 0.02 | 0.0062 | 0.006 | 0.032 | IGHV3-66 | IGKV1-33 |
| **VA14_43** | RBD | 3 | 99 | 0.029 | 0.0044 | 0.0037 | 0.0097 | 0.017 | IGHV3-53 | IGKV1-9 |
| **VA14_26** | RBD | 2 | 98 | 0.21 | 0.17 | 0.093 | 0.11 | 0.11 | IGHV4-34 | IGKV1-5 |
| **VA14_61** | RBD | 4 | 96 | 0.071 | 0.016 | 0.0069 | 0.006 | 0.032 | IGHV1-69 | IGKV4-1 |
| **VA14_33** | RBD | 4 | 90 | 0.056 | 0.031 | 0.0056 | 0.0056 | 0.013 | IGHV3-74 | IGLV6-57 |
| **VA14_36** | RBD | 4 | 72 | 0.025 | 0.02 | 0.013 | 0.0062 | 0.02 | IGHV4-59 | IGKV1-33 |
| **VA14_65** | RBD | 4 | 59 | 0.047 | 0.038 | 0.047 | 0.27 | >50 | IGHV3-21 | IGKV1-39 |
| **VA14_49** | RBD | 4 | 55 | 3.06 | 0.91 | 1.31 | 3.04 | 2.16 | IGHV3-30 | IGKV1-33 |
| **VA14_19** | RBD | 4 | 55 | 0.79 | 0.32 | 0.24 | 1.15 | 0.29 | IGHV3-15 | IGKV1-39 |
| **VA14_R38** | RBD | 4 | 45 | 0.038 | 0.04 | 0.053 | 0.092 | 0.04 | IGHV3-43 | IGKV3-20 |
| **VA14_50** | RBD | 4 | 45 | 0.031 | 0.024 | 0.66 | 1.39 | >50 | IGHV1-18 | IGKV3-11 |
| **VA14_10** | RBD | 4 | 45 | 0.78 | 0.13 | 3.7 | 4.95 | 0.25 | IGHV3-23 | IGKV1-33 |
| **VA14_67** | RBD | 4 | 43 | 0.029 | 0.022 | 0.049 | 0.29 | 0.055 | IGHV5-51 | IGKV1-39 |
| **VA14_07** | RBD | 4 | 43 | 0.28 | 0.19 | 0.46 | 1.16 | 0.86 | IGHV4-39 | IGKV3-20 |
| **VA14_27** | RBD | 4 | 34 | 0.24 | 0.18 | 0.16 | 0.73 | 1.98 | IGHV5-51 | IGKV1-39 |
| **VA14_09** | RBD | 4 | 28 | 0.027 | 0.019 | 11.9 | >50 | >50 | IGHV1-18 | IGKV3-11 |
| **VA14_16** | NTD | 8 | 23 | 0.14 | 0.051 | 0.14 | 0.22 | 0.027 | IGHV3-48 | IGKV1-17 |
| **VA14_68** | NTD | 8 | 19 | 0.39 | 0.075 | 10.19 | 0.48 | 0.067 | IGHV4-59 | IGKV4-1 |
| **VA14_21** | NTD | 5 | 58 | 1.22 | 0.84 | 13.4 | 5.17 | 14.07 | IGHV1-69 | IGKV4-1 |
| **VA14_22** | NTD | 5 | 51 | 0.13 | 0.53 | >50 | 0.78 | >50 | IGHV3-48 | IGKV1D-12 |
| **VA14_58** | NTD | 6 | 28 | >50 | 0.0085 | 0.026 | >50 | >50 | IGHV1-8 | IGKV3-11 |
| **VA14_25** | NTD | 6 | 19 | >50 | 0.052 | 0.093 | >50 | >50 | IGHV1-69 | IGKV3-15 |
| **VA14_47** | S-only | 7 | 59 | 2.1 | >50 | >50 | >50 | >50 | IGHV3-30 | IGKV3-20 |

**Supplementary table S1: Binding and neutralization properties of isolated mAbs.**
